# Non-Invasive Confocal Fluorescence Imaging of Mice Beyond 1700 nm Using Superconducting Nanowire Single-Photon Detectors

**DOI:** 10.1101/2021.08.13.456312

**Authors:** Feifei Wang, Fuqiang Ren, Zhuoran Ma, Liangqiong Qu, Ronan Gourgues, Chun Xu, Ani Baghdasaryan, Jiachen Li, Iman Esmaeil Zadeh, Johannes WN Los, Andreas Fognini, Jessie Qin-Dregely, Hongjie Dai

**Affiliations:** Department of Chemistry and Bio-X, Stanford University, Stanford, CA 94305, USA; School of Medicine, Stanford University, Stanford, CA 94303, USA; Single Quantum B.V., Delft 2628 CJ, The Netherlands; Department of Imaging Physics, Delft University of Technology, Delft 2628 CJ, The Netherlands

## Abstract

Light scattering by biological tissues sets a limit to the penetration depth of high-resolution optical microscopy imaging of live mammals *in vivo*. An effective approach to reduce light scattering and increase imaging depth is by extending the excitation and emission wavelengths to the > 1000 nm second near-infrared (NIR-II), also called the short-wavelength infrared (SWIR) window. Here, we developed biocompatible core-shell lead sulfide/cadmium sulfide (PbS/CdS) quantum dots emitting at ~1880 nm and superconducting nanowire single photon detectors (SNSPD) for single-photon detection up to 2000 nm, enabling one-photon fluorescence imaging window in the 1700-2000 nm (NIR-IIc) range. Confocal fluorescence imaging in NIR-IIc reached an imaging depth of ~ 800 μm through intact mouse head, and enabled non-invasive imaging of inguinal lymph nodes (LNs) without any surgery. In vivo molecular imaging of high endothelial venules (HEVs) with diameter down to ~ 6.6 μm in the lymph nodes was achieved, opening the possibility of non-invasive imaging of immune trafficking in lymph nodes at the single-cell/vessel level longitudinally.

---

In vivo high resolution optical microscopic imaging of mice has empowered investigations of biological structures, molecular nature/identity of cell surface receptors and cellular processes, events and functions at the single-cell level. However, the heterogeneous nature and complex compositions of biological tissues present a major challenge, limiting the imaging penetration and signal to background ratios (SBR) due to light scattering and indigenous tissue autofluorescence. Extending the excitation wavelength to the NIR and SWIR (900-3000 nm) range by multi-photon (MP) imaging has been highly successful in suppressing scattering and affording greater penetration depths, but still relies on invasive surgery to expose underlying organs such as the brain and lymph nodes in order to afford sufficient imaging depths and resolution ^1–6^. Up to 1700 nm excitation has been employed by three-photon microscopy^7^, enabling > 500 μm deep through-skull (with scalp removed) mouse brain 3D volumetric imaging^8^.

In recent years various fluorescent/luminescence dyes and nanoparticle probes with emission in the 1000-1700 nm range have been developed, including small organic molecules^9,10^, carbon nanotubes (CNTs)^11^, quantum dots^12,13^ and rare-earth nanoparticles^14–16^. Using these dyes and probes confocal^12,17^ and light-sheet microscopy^18,19^ employed one-photon excitation up to 1540 nm and emission up to 1700 nm to benefit from reduced scattering of both excitation and emission light^19^, affording non-invasive in vivo microscopy of blood micro-vessels and molecular imaging at single-cell level. In vivo NIR-II microscopy imaging has facilitated investigating mouse models of cardiovascular diseases, brain injury and cancer, as exemplified by imaging of CD4, CD8, OX40 and other bio-markers on immune cells in the tumor microenvironment in response to immunotherapy^12,18–20^.

One of the limiting factors of optical imaging depth into biological tissues is water absorption^21^ that exhibits a local peak at ~ 1445 nm due to the vibrational overtone mode of O-H bond bending (Fig. 1a, and Supplementary Fig. 1 for linear scale absorbance plot). In the 1000-1400 nm range (including the previously defined NIR-IIa sub-window of 1300-1400 nm)^22^ light absorption is relatively low and imaging depths in tissues such as the brain are dominant by scattering (Fig. 1b). Both absorption and scattering influence NIR-IIb (defined as 1500-1700 nm)^23^ imaging depth in mouse brain, with an effective attenuation length^1^ of *l*_e_ = 1/(1/*l*_s_+1/*l*_a_), where *l*_s_ is the scattering length in mouse brain and can be mimicked by 5% intralipid^18^. *l*_a_ is the attenuation length of light due to water absorption (Fig. 1b). The NIR-IIb sub-window affords a longer attenuation length *l*_e_ than in 1000-1400 nm due to reduced light scattering (Fig.1b), allowing for deeper imaging. The 1700 nm NIR-IIb border-line is set by the upper detection limit of indium gallium arsenide (InGaAs) detectors (900-1700 nm) commonly used for NIR-II/SWIR imaging. Beyond 1700 nm, light scattering further reduces but water adsorption increases (Fig.1a,b). Here we define the 1700-2000 nm as the NIR-IIc sub-window and revised the NIR-II window to be 1000-3000 nm (with the upper bound being the same as SWIR^24^). Also, the final sub-window in NIR-II/SWIR for one-photon imaging would be ~ 2100-2300 nm (defined here as the NIR-IId sub-window) range with similar attenuation length as NIR-IIc (Fig. 1b), beyond which water absorption becomes overwhelming and through-tissue fluorescence imaging is impossible (Fig. 1a). Thus far there has been no report of one-photon fluorescence imaging of living systems in the > 1700 nm range, limited by the lack of biocompatible fluorescent/luminescent probes with sufficient quantum yield/brightness, and that InGaAs cameras/detectors are insensitive to light > 1700 nm in the NIR-IIc/NIR-IId range.

**Figure 1.**
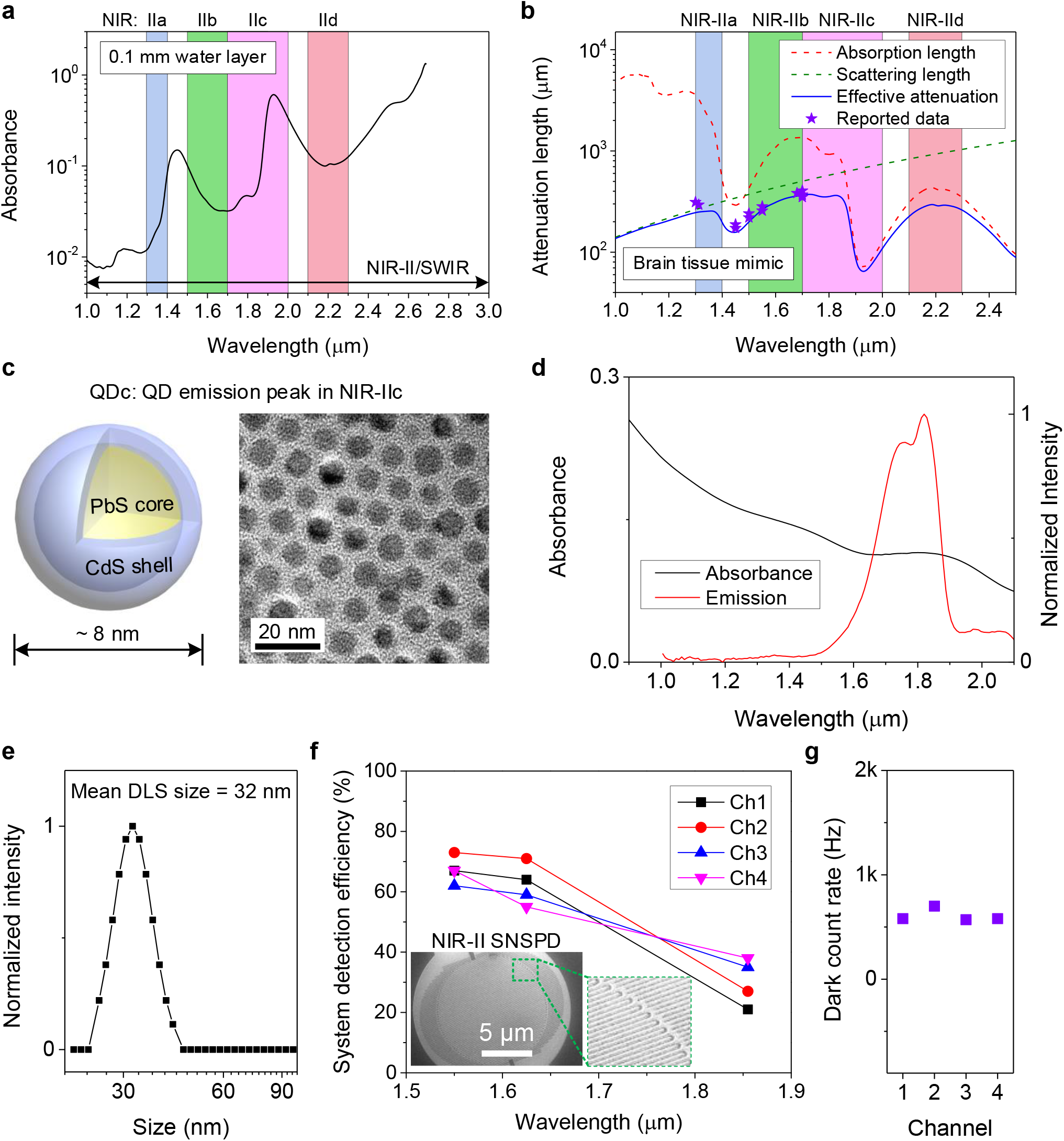
Lead sulfide quantum dots and superconducting nanowire single photon detectors enabling fluorescence imaging beyond 1700 nm. (**a**) The absorbance of water measured in a cuvette with 0.1 mm light path. (**b**) Wavelength-dependent attenuation length (1/(1/*l*_s_+1/*l*_a_)) of brain tissue mimicked by 5% intralipid solution. The scattering length (*l*_s_) of mouse brain was mimicked by 5% intralipid solution^18^. *l*_a_ = 1/*μ*_a_ = *L*/(*D* × ln10) is the attenuation length of light due to water absorption^38^, where *μ*_a_ is absorption coefficient, *L* is optical pathlength and *D* is absorbance as shown in (**a**). The stars represent previously reported effective attenuation length of mouse brain measured in vivo^1,39^. (**c**) Schematic design of NIR-IIc core-shell PbS/CdS quantum dots (left) and corresponding transmission electron microscopy image (right). (**d**) The absorption spectrum of QDc and emission spectra of P^3^-QDc measured in tetrachloroethylene (TCE) or PBS buffer, respectively. A 1-mm cuvette was used. The absorption spectrum was measured in organic phase as the serious absorption of water at ~1445 nm influenced the measurement of absorbance in water. (**e**) Dynamic light scattering spectra of P^3^-QDc with polymeric crosslinked network in PBS buffer. (**f**) Optimized multi-channel SNSPD with high system detection efficiency in NIR-IIb (Ch-1and Ch-2) and NIR-IIc (Ch-3and Ch-4) windows. Insets: A scanning electron microscopy image of a typical SNSPD. (**g**) The dark counts of four SNSPD used in our experiments.

Here we developed aqueous soluble core-shell lead sulfide/cadmium sulfide quantum dots (PbS/CdS QD) with emission peak ~ 1880 nm and employed superconducting nanowire single photon detectors (SNSPD) for in vivo imaging in the > 1700 nm range to further suppress light scattering and push the limit of one-photon through-tissue imaging. Confocal microscopy through mouse tissues in the NIR-IIc window revealed mouse scalp/skull/brain structures in 3D and enabled surgery-free molecular imaging of the peripheral node addressin (PNAd) on HEVs inside lymph nodes non-invasively.

We synthesized PbS quantum dots with emission peak at ~ 2009 nm via a modified organometallic route^25^. A CdS shell on the PbS core was then grown by the cation exchange approach to protect the PbS core from degradation^12^. The shell growth step shifted the emission peak of the core to ~ 1880 nm (Supplementary Fig. 2a, see synthesis procedure in Methods). The resulting PbS/CdS QD with emission peak in NIR-IIc (named ‘QDc’ herein) showed a narrow size distribution around ~ 8 nm (Fig. 1c and Supplementary Fig. 2b), larger than our previous ~ 6.9 nm QD with emission peak in NIR-IIb (named ‘QDb’)^12^. To transfer the QDc from organic phase to aqueous phase, a hydrophilic polymeric cross-linked network (P^3^ coating) developed by our group was coated on the QDc to impart biocompatibility in physiological environments and biliary excretion in ~ 2 weeks without apparent toxic effects^14,26^. The final aqueous stabilized P^3^-QDc exhibited a peak emission wavelength of ~ 1820 nm in NIR-IIc (Fig. 1d). Dynamic light scattering (DLS) analysis of the P^3^-QDc showed an average hydrodynamic size of ~ 32 nm in aqueous solutions (Fig. 1e).

The upper spectral detection limit of 1700 nm for InGaAs detectors was overcome by SNSPD designed for wavelengths from 1550 nm to 2000 nm. The single-photon detectors were fabricated on an optimized niobium titanium nitride (NbTiN) superconducting film^27^ with an optimized superconducting nanowire width and detector geometry. In this system, detectors with cavities based on an Au back mirror with a SiO_2_ *λ*/4 (*λ* is the wavelength) spacer between the mirror and the superconducting film, and cavities based on a Distributed Bragg Reflector (DBR) were constructed^28^. These two types of detectors with different cavities performed similarly in terms of efficiency and dark-counts in the wavelength range of 1550-2000 nm. The specially designed SNSPD showed a system detection efficiency (SDE) of ~ 50% to 75% in the NIR-IIb 1500-1700 nm window (Channel 1 and 2) and ~ 25% to 50% in the NIR-IIc 1700-2000 nm window (Channel 3 and 4), respectively (Fig. 1f). These SNSPD were cooled at ~ 3 K Gifford-McMahon Cryocooler. The SNSPD were connected to our home-built confocal microscope through single mode (SM) fibers (~ 10 μm mode field diameter) transmitting in 1200-2000 nm (see Method and Supplementary Figs. 3 and 4) that also functioned as a pinhole to reject the out-of-focus fluorescence signal. The home-built confocal microscope allowed diffraction-limited resolution in non-scattering media calibrated by observing nanoparticles on a cover glass (see resolution analysis in Methods). The selected SM fiber afforded a natural cut-off for the black-body radiation that significantly reduced the dark-count rate to be ~ 500 Hz and enabled high SBR for NIR-IIc fluorescence imaging (Fig. 1g and Methods). The low timing jitter (< 55 ps) facilitated high temporal single-photon resolution. Note that a previous work employed SNSPD with SDE > 50% at 1064 nm for cerebral vasculature (labeled by the IR820 dyes) imaging through skull with scalp removed in the ~ 1000-1200 nm detection range^29^.

We first imaged a 50 μm diameter capillary filled with aqueous suspensions of P^3^-QDb or P^3^-QDc with emission peak residing in NIR-IIb (1500-1700 nm)^12^ or NIR-IIc (1700-2000 nm) immersed in ~ 5% intralipid solution, mimicking blood vessels in a mouse brain tissue (Fig. 2a-d). NIR-IIc fluorescence was collected in 1750-2000 nm (see Method for filters used), but at millimeters imaging depths the actual upper limit was ~ 1900 nm set by water absorption (Fig.1a). This was also gleaned by that the detected fluorescence signals of QDc through 2-10 mm thick layers of water showed diminished > 1900 nm signals due to water absorption (Fig. 2e), while emission was clearly detected in the 1600-1850 nm range even through a 10 mm thick water layer. For wide-field imaging of QDb filled capillary in the 1580-1700 nm range of NIR-IIb with an InGaAs camera (excitation 808 nm), the capillary was well resolved at up to ~ 2.3 mm immersion depth in the intralipid solution but not at 2.6 mm depth (Fig. 2a, left and Fig. 2b). Quasi confocal microscopy (with a 5X objective, *z* resolution ~ 150 μm) imaging in the same 1580-1700 nm range using the SNSPD (under 1540 nm laser excitation) afforded improved SBR over InGaAs camera imaging (Fig. 2d), but was still incapable of resolving the capillary beyond ~ 2.3 mm depth (Fig. 2a, middle). The ‘sidelobes’ around the capillary in the wide-field images were attributed to scattered light and can be removed by the confocal microscopy (Fig. 2a). In contrast, confocal microscopy (1750-2000 nm emission; 1540 nm excitation) imaging of QDc filled capillary in NIR-IIc using the SNSPD extended the imaging depth to ~ 2.6 mm (Fig. 2a, right) and afforded higher SBR than NIR-IIb imaging at all depths (Fig. 2d).

**Figure 2.**
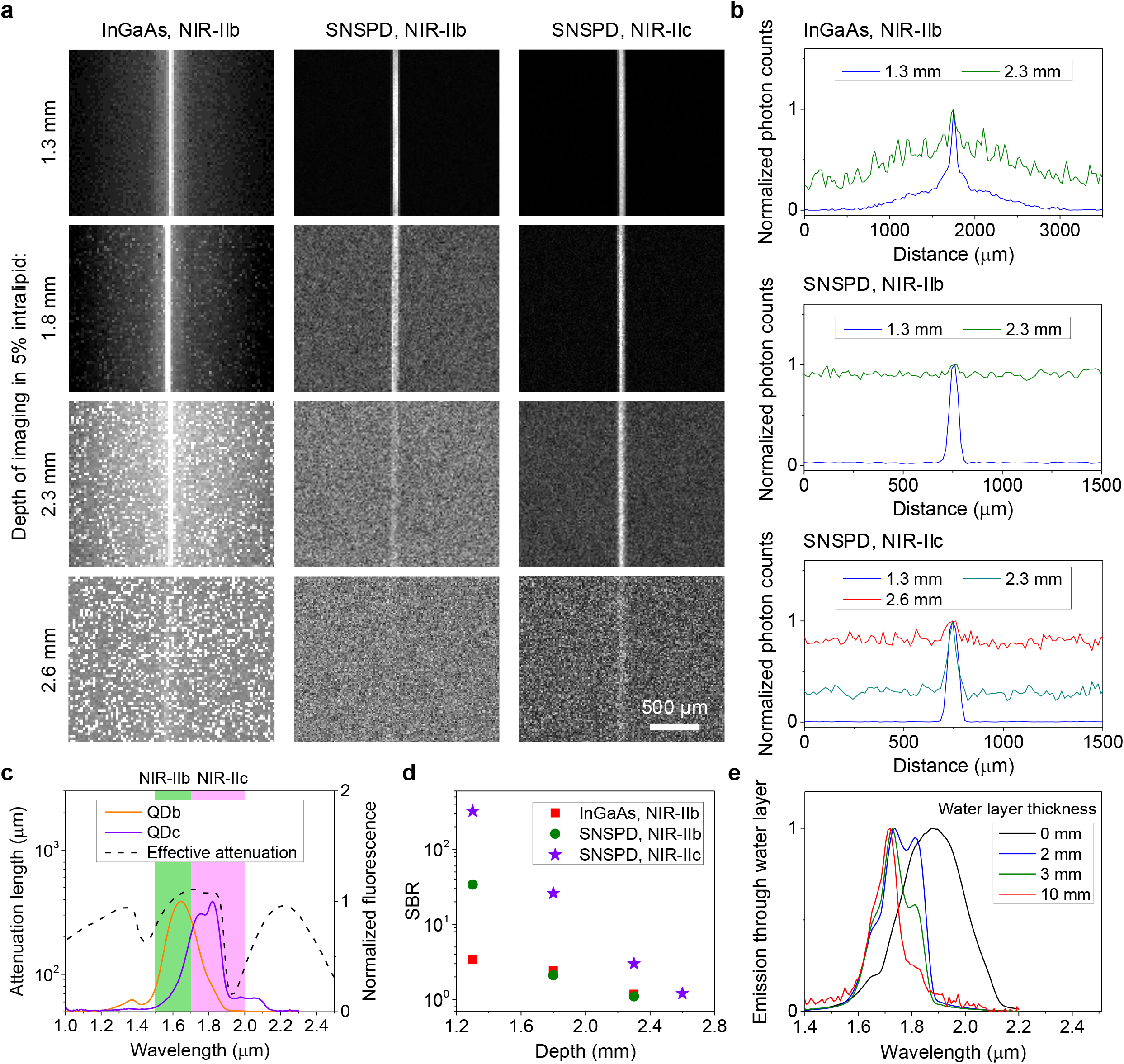
Fluorescence imaging in NIR-IIb and NIR-IIc windows. (**a**) Fluorescence imaging of a 50-μm diameter capillary tube filled with NIR-IIb QDb or NIR-IIc QDc immersed at different depths in 5% intralipid by a wide-field system with a 2D InGaAs camera or a confocal microscope with SNSPDs. The 50-μm diameter capillary tubes immersed in 5% intralipid solution mimicked blood vessels in mouse brain tissues. An 808-nm laser was used for NIR-IIb wide-field imaging. A 1540-nm laser and a 5X objective (NA = 0.12) were used for NIR-IIb and NIR-IIc confocal microscopy. NIR-IIb and NIR-IIc fluorescence was collected in 1580-1700 nm or 1750-2000 nm ranges, respectively. (**b**) Normalized photon counts profiles of the capillary immersed at different depth in the 5% intralipid imaged by a wide-field system or a confocal microscope in NIR-IIb or NIR-IIc window. (**c**) Fluorescence spectra of P^3^-QDb and P^3^-QDc in PBS with emission peaks in NIR-IIb^12^ and NIR-IIc regions respectively. A cuvette with 1-mm light path was used to measure the spectrum. The effective attenuation length is from Fig. 1b. (**d**) Comparison of SBR of the results shown in (**a**). The SBR of wide-field results was from the ratio of the capillary brightness and the signal from sidelobe around the capillary. (**e**) Emission spectra of NIR-IIc QDc in a cuvette measured under water at depths of 2 mm, 3 mm and 10 mm respectively. The 0 mm data indicates no water layer and the emission spectrum was measured with QDc in tetrachloroethylene without any influence from water.

We also performed confocal imaging using a 1319 nm laser for excitation and the same SNSPD. Compared to 1540 nm excitation, the maximum imaging depth decreased from ~ 2.3 mm to ~ 1.8 mm and ~ 2.6 mm to ~ 2.3 mm for NIR-IIb and NIR-IIc confocal microscopy respectively (Supplementary Fig. 5). Together, these results confirmed that longer excitation and emission wavelength allowed for deeper imaging penetration depth in scattering media.

Upon tail-vein injection of P^3^-QDc, we performed NIR-IIc non-invasive 3D confocal imaging of blood vessels through intact mouse head (Fig. 3a,b) in 1.1 μm and 5 μm steps scanned in *xy*-plane and *z*-axis respectively, with per-plane imaging time linearly increased from 5 s to 30 s along *z* to compensate for weaker signals at greater depths due to tissue scattering/absorption. The P^3^-QDc circulating in the blood vessels in intact mouse head layers including scalp, skull, meninges and ~ 550 μm thick brain cortex were clearly resolved, with a 3D volumetric imaging depth of ~ 800 μm through the intact mouse head (Fig. 3b and Supplementary Video 1 for 3D views). Vessel like channels through the meninges connecting the skull and brain cortex were observed (Fig.3b, pointed by arrows). These vessels/channels were shown important to immune responses of the brain to injuries and infections^18,30^.

**Figure 3.**
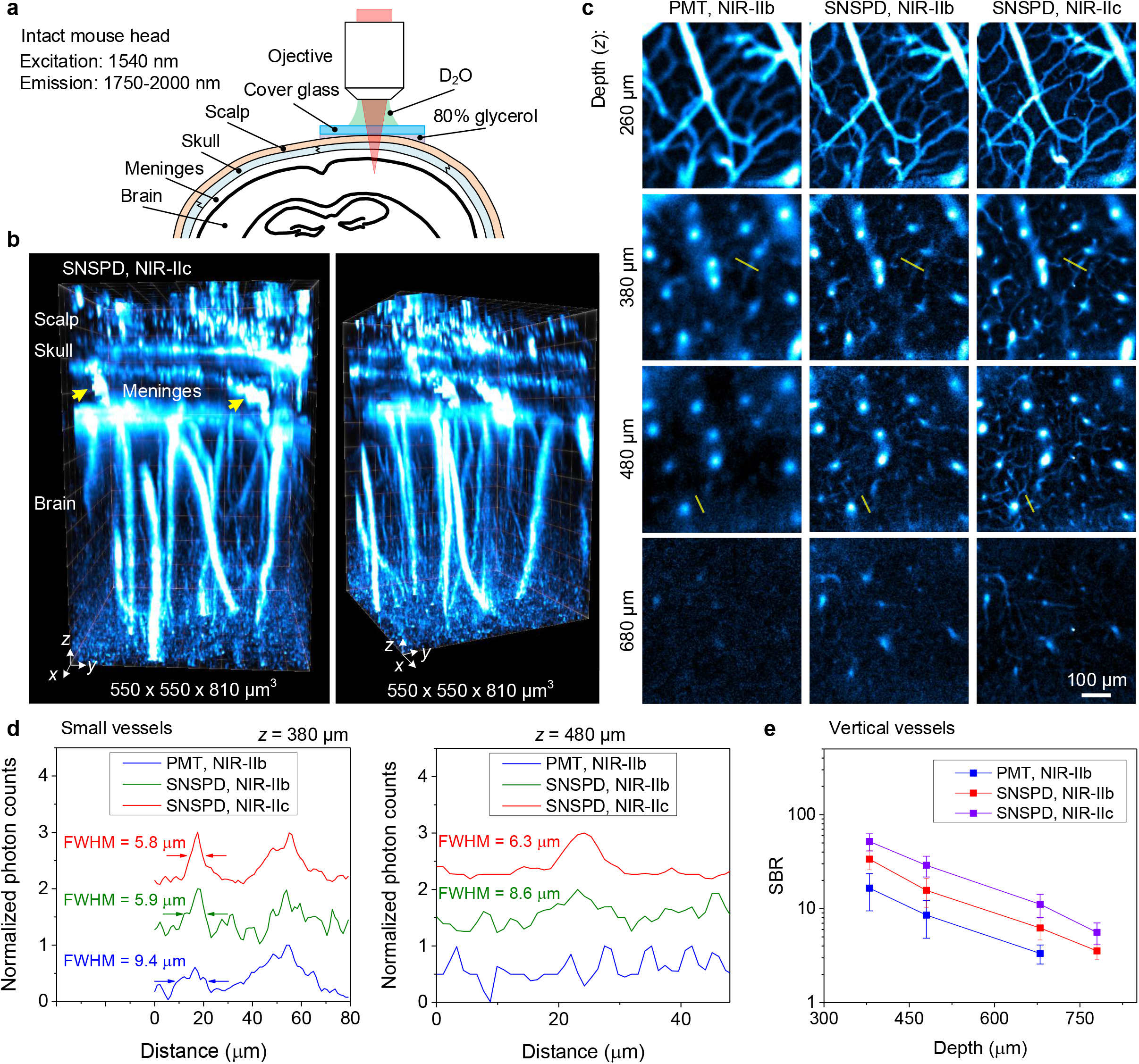
Non-invasive *in vivo* confocal microscopy of intact mouse head in NIR-IIc window. (**a**) Schematic of intact mouse head imaging in NIR-IIc window. The gap between a ~170-μm thick cover glass and mouse head were filled with 80% glycerol. A 25X objective (NA = 1.05) and heavy water (D_2_O) as the immersion liquid was used. (**b**) 3D volumetric images of blood vessels in an intact mouse head visualized through the scalp, skull, meninges and brain cortex obtained with a 5-μm *z*-scan increment. The imaging time per plane was linearly increased from 5 s to 30 s from the first frame at the scalp surface to the last plane inside the brain tissue. The arrows point to the vessel like channels connecting the skull and brain cortex in the meninges. Confocal microscopy was performed 30 min after intravenous injection of P^3^-QDc. Fluorescence signal was collected in 1750-2000 nm window and excited by a 1540 nm laser. (**c**) High-resolution confocal images of blood vessels at various depths through intact mouse head imaged with a PMT or SNSPD in NIR-IIb or NIR-IIc window. Imaging was performed 30 min after intravenous injection of P^3^-QDb and P^3^-QDc sequentially. NIR-IIb and NIR-IIc fluorescence was collected in 1500-1700 nm and 1750-2000 nm windows and excited by a 1319-nm and a 1540-nm laser, respectively. (**d**) Normalized photon counts profiles along the yellow lines in **c**. (**e**) Comparison of SBR for *xy* images recorded at different depths by confocal microscopy with a PMT or SNSPD. The signals of vertical vessels were used to calculate the SBR as these vessels existed from the surface of brain cortex to the deepest layer of imaging. Data are shown as mean ± s.d. derived from analyzing > 5 vessels at each depth.

We compared SNSPD to a commercial 900-1700 nm photomultiplier tube (PMT) for non-invasive confocal microscopy imaging through intact mouse head. For this a mouse was intravenously injected with P^3^-QDb and P^3^-QDc sequentially 30 min before imaging (see Method). For NIR-IIb imaging under the same 1319 nm laser excitation, the SNSPD based confocal microscopy resolved small capillary vessels not detected by PMT and allowed higher spatial resolution at greater imaging depths (Fig. 3c left column for PMT vs. middle for SNSPD, and Fig. 3d). Under a laser excitation of 1540 nm, imaging in the NIR-IIc window afforded higher resolution than in the NIR-IIb region under an excitation of 1319 nm, using the same SNSPD (Fig. 3c, middle, right and Fig. 3d). For example, the full width at half maximum (FWHM) of a vessel imaged at *z* = 380 μm was measured to be 9.4 μm, 5.9 μm and 5.8 μm by confocal microscopy using PMT in NIR-IIb, SNSPD in NIR-IIb and NIR-IIc, respectively (Fig. 3d left). As the imaging depth increased to 480 μm, vessels not resolved by PMT were resolved in NIR-IIb and NIR-IIc widows (FWHM of 8.6 μm and 6.3 μm respectively in Fig.3d) using SNSPD. Confocal microscopy in NIR-IIc window afforded the highest contrast, with a SBR of ~ 50 at z = 380 μm that decreased to ~ 6 at 780 μm, which was ~ 1.7 times and ~ 3.3 times higher than NIR-IIb imaging recorded by SNSPD and PMT respectively (Fig. 3e).

Next we performed non-invasive in vivo NIR-IIc confocal microscopy imaging (excitation 1540 nm, emission 1750-2000 nm) of mouse lymph nodes (LNs) without performing any surgery to expose the LNs (Fig. 4a). The lymphatic system plays crucial roles in immune responses to infections, cancer and vaccination. It is highly desirable to image and monitor cellular immunological events within the lymph nodes in real-time and over-time longitudinally with cellular resolution. Here we focused on imaging high endothelial venules (HEVs) in LNs that are small postcapillary venules responsible for mediating entry of immune cells from blood circulation into LNs^31^. In cancer, sentinel LN HEVs are remodeled before metastatic tumor cells are observed in the LNs^32^. Recent work also revealed HEVs providing an exit route for tumor cell dissemination and metastases^33,34^. For these reasons in imaging of sentinel LN HEVs could lead to new insights into immune responses to various infectious diseases and cancer^31^. Thus far in vivo visualization of LN HEVs has been limited to intravital microscopy with installed transparent windows on mice via invasive surgery^33^. For longitudinal observation of HEVs in experiments monitoring metastatic tumor cells invading HEVs, LNs had to be harvested at serial timepoints from different mice and imaged ex vivo^32,34^.

**Figure 4.**
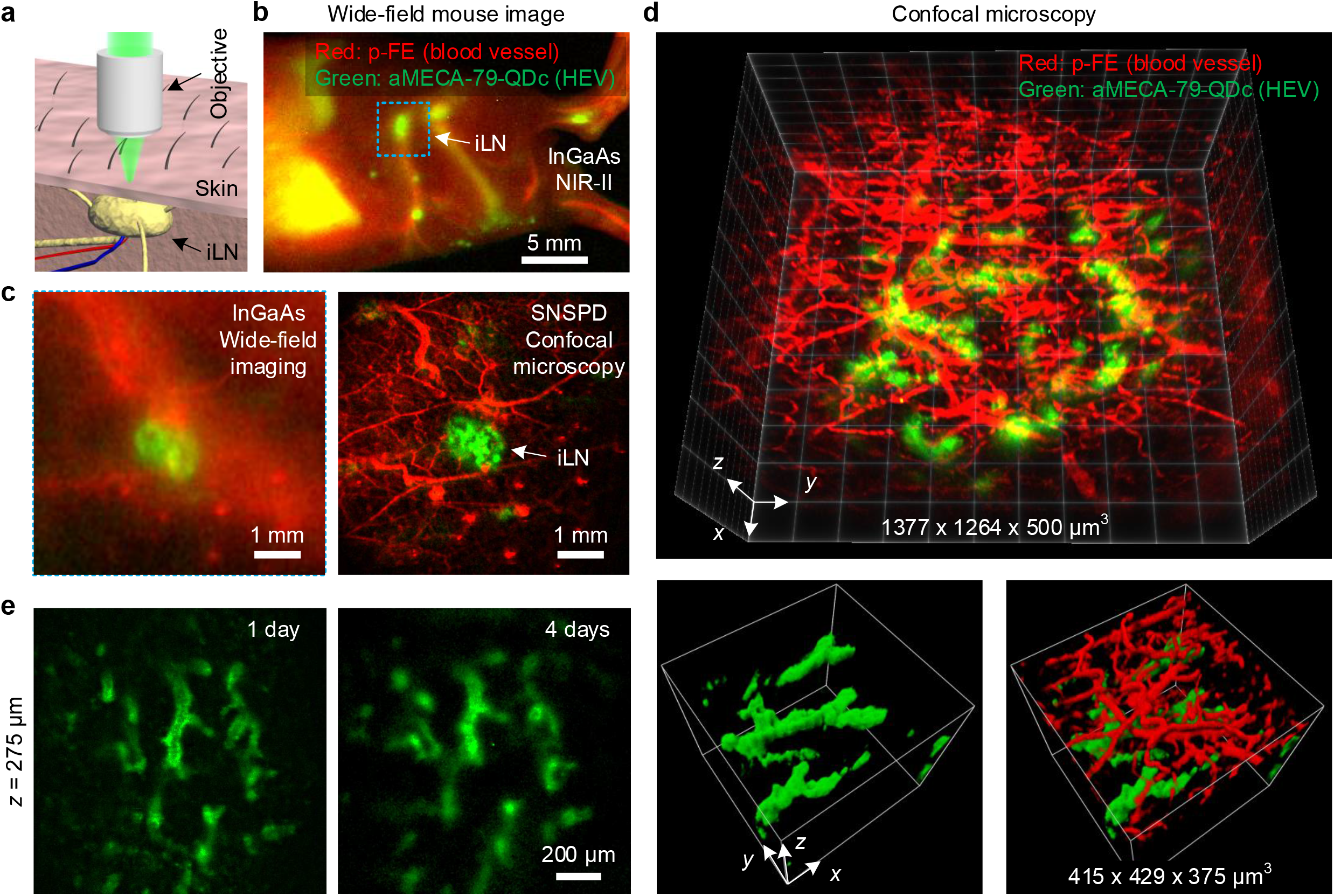
Non-invasive *in vivo* NIR-II confocal microscopy of inguinal lymph node in mice. (**a**) Schematic of non-invasive imaging of iLN by confocal microscope. (**b**) Wide-field imaging of p-FE labelled blood vessels (red, excitation: 808 nm, emission: 1200-1400 nm, exposure: 30 ms) and aMECA-79-QDc labelled HEVs (green, excitation: 808 nm, emission: 1500-1700 nm, exposure: 40 ms) 1 day post intravenous injection of aMECA-79-QDc and 10 min after intravenous injection of free p-FE. (**c**) Comparison of wide-field imaging and confocal microscopy of inguinal lymph node marked by the rectangular in (**b**) at a higher magnification. An 808 nm laser was used for wide-field imaging to excite p-FE (emission: 1200-1400 nm, exposure: 40 ms) and aMECA-79-QDc (emission: 1500-1700 nm, exposure: 40 ms). For confocal microscopy, p-FE (emission: 1200-1400 nm, imaging time: 25s) and aMECA-79-QDc (emission: 1750-2000 nm, imaging time: 25s) were excited by an 808 nm laser and a 1540 nm laser, respectively. A 5X objective was used. (**d**) High-resolution confocal microscopy of blood vessels (red, p-FE dye circulating in blood vessels; excitation: 808 nm, emission: 1200-1400 nm) and HEVs labelled by aMECA-79-QDc (green, excitation: 1540 nm, emission: 1750-2000 nm) in iLN with a 5-μm *z*-scan increment. The imaging time per plane was linearly increased from 5 s to 15 s from the first frame at the skin surface to the last plane inside the iLN. Skin layer vessels (red) and underlying HEVs (green) in iLN were resolved. (**e**) Longitudinal observation of aMECA-79-QDc labelled HEVs in iLN 1 day and 4 days post intravenous injection. To image the same iLN multiple times, we first located the iLN by wide-field imaging of aMECA-79-QDc. We then focused onto the iLN for confocal microscopy as only the iLN showed strong fluorescence signal due to the antibody-QDc labeled HEVs. (**d**,**e**) A 25X objective (NA = 1.05) was used.

We performed non-invasive in vivo NIR-IIc molecular imaging of the PNAd on HEVs in inguinal lymph nodes (iLNs) of mice (Fig. 4). We conjugated anti-MECA-79 antibodies to P^3^-QDc (aMECA-79-QDc), intravenously injected into a mouse through the tail-vein, and 24 h later injected an organic probe p-FE (excitation: 808 nm; emission: 1200-1400 nm)^20^ as a blood vessel NIR-II imaging agent. The MECA-79 antibody was known to specifically target the 6-sulfo sialyl Lewis X epitope on PNAd expressed on HEV^32^. Ten minutes after injection of p-FE, we first performed wide-field imaging in NIR-IIb (the QDc exhibited emission in both NIR-IIb and NIR-IIc) under an excitation of 808 nm (detection 1500-1700 nm) and observed anti-MECA-79 targeted QDc emission in the iLNs, with blood vessels labeled by circulating p-FE imaged in the 1200-1400 nm emission range (Fig. 4b, Fig. 4c left). In comparison, no QDc emission signal was detected in the iLNs in a control mouse injected with free P^3^-QDc without anti-MECA-79 conjugation (Supplementary Fig. 6). The results confirmed specific molecular targeting of HEVs in LNs with aMECA-79-QDc injected intravenously.

We then performed non-invasive two-color confocal microscopy imaging to map out/differentiate the spatial distributions of blood vessels (with circulating p-FE dye) and HEVs (aMECA-79-QDc) in the iLN. At low magnifications the iLN region was imaged by both InGaAs wide-field system and SNSPD confocal microscopy (Fig. 4c). The latter clearly rejected scattering induced background and allowed weak signal detection in both 1200-1400 nm and NIR-IIc window. High-resolution 3D confocal imaging of iLNs revealed HEVs in the iLNs below a skin layer with p-FE labeling the skin blood vessels (Fig. 4d). Confocal imaging at different depths showed that only HEVs were labelled by aMECA-79-QDc, while both HEVs and the surrounding blood vessels in the iLN were labeled by p-FE (Supplementary Fig. 7), confirming specific targeting of HEVs by aMECA-79-QDc. Confocal microscopy in NIR-IIc allowed iLNs molecular imaging with a penetration depth of ~ 500 μm including the skin layer, which was ~ 2 X deeper than in imaging in the 1200-1400 nm range (Supplementary Fig. 7). The high stability of P^3^-QDc and the aMECA-79-QDc afforded lasting targeting effect, allowing for non-invasive NIR-IIc confocal microscopy imaging of HEVs in iLNs longitudinally over more than 4 days (Fig. 4e).

In this work, we developed biocompatible PbS/CdS QD emitting at ~ 1880 nm and employed SNSPD for sensitive 1550-2000 nm photon detection, opening a new NIR-IIc sub-window (1700-2000 nm) for one-photon fluorescence imaging under laser excitations up to 1540 nm, suppressing scattering and enhancing penetration depth for both excitation and emission in the 1000-3000 nm NIR-II/SWIR window. NIR-IIc confocal microscopy with single photon detectors afforded non-invasive high-resolution imaging of intact mouse head and iLNs with penetration depths of ~ 800 μm and ~ 500 μm, respectively. Imaging could be further advanced by extending the excitation to > 1700 nm and emission to the NIR-IId (2100-2300 nm) sub-window, which would further increase in vivo microscopy imaging depth if the challenge to making bright probes in this sub-window of NIR-II/SWIR can be met. Image scanning microscopy^35^ and the measurement of quantum photon correlation^36^ could be introduced to further improve the resolution of NIR-IIc confocal microscopy. Non-invasive, deep penetrating 3D microscopy imaging has the potential to emerge as a powerful tool for longitudinal monitoring the immune responses and cellular functions and events in lymph nodes, such as the entry of T cells from blood circulation into LNs to interact with antigen-presenting cells for T cell activation in responses to infections or immunotherapy^31,32^.

## Methods

### Materials

Lead (II) chloride (PbCl_2_), 1-Octadecene (ODE), cadmium oxide (CdO) and sulfur powder (sublimed) were purchased from Alfa Aesar. Oleylamine, oleic acid, 1-octadecene, poly(maleic anhydride-alt-1-octadecene) (PMH; average molecular weight: 30k-50k), 4-morpholineethanesulfonic acid (MES), 4-(dimethylamino)pyridine (DMAP), poly(acrylic acid) (PAA; average molecular weight: 1800), 1-(3-dimethylaminopropyl)-3-ethylcarbodiimide hydrochloride (EDC), and 2-Amino-2-(hydroxymethyl)-1,3-propanediol (tris-base) were purchased from Sigma-Aldrich. Hexane, toluene, chloroform, and DI water were purchased from Fisher Scientific. Methoxy polyethylene glycol amine (mPEG-NH2; average molecular weight: 5 kD) was purchased from Laysan-Bio. 8-arm polyethylene glycol amine (8Arm-PEG-NH_2_·HCl; average molecular weight: 40 kDa) was purchased from Advanced Biochemicals. All the chemicals were used without further purification. 50 μm capillaries were bought from VitroCom. Purified anti-mouse/human PNAd antibody (Clone: MECA-79) was purchased from BioLegend.

### Synthesis of NIR-IIc PbS/CdS QD

The synthesis of NIR-IIb PbS/CdS QDb can be found in Refence 12. This work developed the synthesis of NIR-IIc PbS/CdS QDc.

The core PbS QD were synthesized using a modified organometallic method^12,25^. In a typical reaction, sulfur precursor solution was prepared by mixing 0.08 g (5 mmol) of sulfur powder, and 7.5 mL of oleylamine in a two-neck flask at 120 °C under argon for 30 min.

Synthesis of PbS quantum dots. 1.668 g (3 mmol) of PbCl_2_ and 15 mL of oleylamine were mixed in a three-neck flask and degassed for 30 min at 120 °C. The solution was then heated to 165 °C under argon, and kept at that temperature for 15 min. 4.5 mL of the sulfur precursor solution (1.5 mmol of S) was injected into the Pb precursor solution (6 mmol of Pb) under stirring. The temperature was maintained at 165 °C throughout the reaction.

After 180 min, the reaction was quenched by adding 20 mL of cold hexane and 30 mL ethanol. The products were collected by centrifugation and re-suspended in a mixture of 15 mL hexane and 30 mL oleic acid. The quantum dots were precipitated via centrifugation after agitated for 10 min. This precipitation procedure with hexane and oleic acid was repeated 3 times. The quantum dots were re-suspended in 20 mL toluene after centrifugation.

Synthesis CdS shell on PbS quantum dots. CdO (1.2 g, 9.2 mmol), oleic acid (8 mL), and ODE (20 mL) were mixed in a tree-neck flask and heated to 200 °C for 1.5 hours under argon. The solution was then cooled down to 100 °C and degassed under vacuum for 30 min to afford a Cd precursor solution. 5 mL of the prepared PbS QD suspended in toluene was bubbled with argon for 10 min, and then injected into the Cd precursor solution. The reaction flask was quenched with 5 mL cold hexane after the growth reaction was conducted at 100 °C for 60 min. The PbS/CdS quantum dots were precipitated with ethanol and then re-dispersed in hexane.

### Surface modification of PbS/CdS QD with P^3^ coating

0.5 mL chloroform solution containing 10 mg PMH was mixed with 2 mg QD dispersed in 0.5 mL cyclohexane. After overnight stirring, the organic solvent was evaporated for 30min by rotary evaporator. The excess residual was heated at 60 °C for 3 hours to fully remove the remaining organic solvent. Afterwards, 1 mL DMAP (10 mg) aqueous solution was added, and the flask was placed in a sonication bath for 10 min to allow the QD to be fully transferred into the water phase. Furthermore, the QD solution was centrifuged at 50000 r.p.m for 2.5 hours. The precipitated were resuspended in 0.5 mL MES (10 mM, pH = 8.5) followed by the addition of 1.5 mg 8Arm-PEG-NH_2_·HCl (MW 40k) in 1 mL MES (pH=8.5) and 1 mg EDC. The solution was reacted for 3 hours on orbital shaker. The remaining carboxylic groups were quenched by adding 5 mg Tris-base and 2.5 mg EDC to the above solution. After completion of the reaction for 3 hours, the product was dialyzed against water for 12 hours (300 kDa) to completely remove any by-products. The purification was completed by washing the supernatant with a centrifuge filter tube (100 kDa) and the concentrated sample inside the filter was dispersed in 0.5 mL MES (10 mM, pH = 8.5) solution. 0.5 mg PAA (MW 1800) in 1 ml MES (10 mM, pH = 8.5) together with 1 mg EDC was added and the QD solution was allowed to react for 1 hour. The possible large floccules were removed by the centrifugation of solution at 4400 r.p.m for 30 min, followed by the washing of supernatant by a centrifuge tube (100 kDa) for 4 times. The as-prepared QD@PMH-8ArmPEG-PAA were re-dispersed in 0.5 mL MES solution for final layer coating. As such, the above QD@PMH-8ArmPEG-PAA were mixed with 0.5 mg mPEG-NH2 (MW 5k), and 0.1 mg 8Arm-PEG-NH_2_·HCl (MW 40k) in 1 mL MES solution (10 mM, pH = 8.5). After adding of 1 mg EDC, and the solution was allowed to react for 3 hours. Then, 5 mg Tris-base and 2.5 mg EDC were added into the above solution and left to react for another 3 hours. The reaction product was centrifuged at 4400 r.p.m for 30 min to remove potential large aggregates, then the supernatant was washed by a centrifuge filter (100 kDa) for 4 times. The final QD@PMH-8ArmPEG-PAA-mixed PEG were dispersed in 500 μl 1xPBS solution for further use.

### Conjugation of MECA-79 antibody on P^3^-QDc (aMECA-79-QDc)

100-μg MECA-79 antibody was washed by centrifuge filter (10 kDa) for 4 times to remove sodium azide. P^3^-QDc (0.25 mg) dispersed in 50 μL 1xPBS solution, MECA-79 antibody (100 μg), EDC (0.6 mg) and 450-μL MES solution (10 mM, pH = 8.5) were mixed and shaken for 3 h. The solution was first centrifuged at 4,400 r.p.m for 30 min to remove potential large floccules. The supernatant was washed by centrifuge filter (10kDa) four times, and then dispersed in 200 μl 1xPBS solution for further injection.

### NIR-II confocal microscope

A laser beam with wavelength of 808 nm, 1319 nm or 1540 nm was reflected by a galvo mirror (GVS002, Thorlabs) into a Plössl scan lens (constructed from two achromatic doublets, L3, L4, AC508-150-C, Thorlabs), a Plössl tube lens (constructed from two achromatic doublets, L1, L2, AC508-300-C, Thorlabs) and then focused by an objective into a sample (Supplementary Fig. 3). Fluorescence signal was collected by the same objective, tube lens and scan lens. After de-scanned by the galvo mirror and resized by two achromatic doublets (L5, AC254-100-C and L6, AC508-150-C), the fluorescence light was focused into three single mode (SM) fibers (P1-SMF28E-FC-5, Thorlabs) by achromatic doublets (L7-L9, AC254-030-C, Thorlabs) after filtered by selected emission filters. Then the SM fiber transmitted the fluorescence to NIR-IIb or NIR-IIc SNSPD. The datasheet of P1-SMF28E-FC-5 only gives attenuation data in 1260-1625 nm. We examined the attenuation of this SM fiber in 1000-2000 nm by measuring the emission spectrum of a mixture of three PbS/CdS QDs (emission peaks at 1100 nm, 1340 nm, 1650 nm and 1880 nm respectively) using a spectrometer connected to a 1D extended InGaAs camera (Princeton Instruments) or SNSPD coupled with the SM fiber. The experimental results show the fiber coupled SNSPD has some attenuation as the wavelength longer than 1678 nm, but the attenuation degree is still acceptable for 1200-2000 nm imaging (Supplementary Fig. 4). These SM fibers showed a natural cut-off of long wavelengths of black-body radiation to reduce the dark-count rate (~ 500 counts per second after fiber coupling). For example, with a similar system equipped with PM2000 fibers (operating wavelength: 1700-2300 nm, Thorlabs) we measured ~10,000 dark-counts per second. The SNSPD were connected to 8-channel electronic driver and mounted in a compact closed-cycle cryostat cooled by a Gifford-McMahon Cryocooler. The temperature of cryostat was stabled as ~ 3 K. The system detection efficiency of the SNSPD was calculated by:

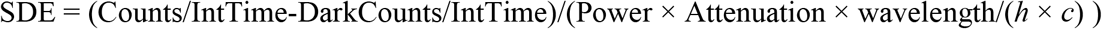

where IntTime is integration time, *h* is Planck constant and *c* is speed of light in free space.

For confocal microscopy with PMT (H12397-75, Hamamatsu), a multimode fiber (M122L01, Thorlabs) with core diameter of 200 μm was used to guarantee enough fluorescence signal detected by the PMT.

The galvo mirror, SNSPD and a motorized translation stage (M-VP-25XL, Newport) were synchronized by an acquisition card (NI PCIe-6374) through Labview. This acquisition card also worked as a photon counter. A 5X objective (NA = 0.12, Leica N Plan), a 10X objective (NA = 0.25, Olympus ULWD MIRPlan), a 20X objective (NA = 0.4, Olympus ULWD MIRPlan), a 20X objective (NA = 0.75, Nikon Plan APO), a 25X objective (NA = 1.05, Olympus XLPLN25XWMP2) and a 100X objective (NA = 0.8, Olympus ULWD MIRPlan) were used in this work. We used heavy water (D_2_O) as the immersion liquid for the 25X objective (NA = 1.05) to minimize the absorption by this immersion layer.

For NIR-IIb imaging, longpass filters with a cut-on wavelength of 1500 nm (FELH1500, Thorlabs) or 1580 nm (BLP01-1550R-25) and a bandpass filter (BBP-1530-1730 nm, Spectrogon) were used to generate imaging windows of ~1500-1700 nm or 1580-1700 nm, respectively. For NIR-IIc imaging, two bandpass filters (FB1750-500 with a transmittance window in 1500-2000 nm and FB2000-500 with a transmittance window in 1750-2250 nm, Thorlabs) were combined for 1750-2000 nm fluorescence collection. A shortpass filter with a cut-off wavelength of 1600 nm (#84-656, Edmund optics) was used to filter 1540 nm laser.

For in vivo imaging, the actual excitation power was approximately 0.15 J cm^−2^, 0.52 J cm^−2^ and 0.95 J cm^−2^ for 808-nm, 1319-nm and 1540-nm lasers, respectively, which is below the safety limit for laser dwell of 114 μs (808 nm, 0.19 J cm^−2^; 1319 nm, 0.57 J cm^−2^; 1540 nm, 1.0 J cm^−2^) as described in Reference ^37^. Excitation power was measured by a laser power meter (3A-SH, NOVA II, OPHIR). Different excitations could be selected by removable mirrors.

### Resolution analysis

We analyzed the spatial resolution of confocal microscopy with SNSPD by experimental measurements and modeling (see Supplementary Note 1). To estimate the system’s best resolution without worsening by scattering, we imaged 300-nm polystyrene beads containing a NIR-II organic dye^20^ with different objectives. The nanoparticles were deposited on a coverslip. Fluorescence emission > 1100 nm was collected at 658-nm excitation. The experimental measured FWHM of nanoparticles were consistent well with the FWHM of theoretically calculated point spread function (PSF) along lateral and vertical directions (Supplementary Fig. 8). When a 25X objective (numerical aperture, NA = 1.05) was used, ~ 0.4 μm in lateral and ~ 1 μm in vertical can be achieved. The agreement of experimental data and modeling suggested validity of the analysis for imaging with the organic dyed beads emitting in the 1000-1300 nm range. Due to the lack of beads emitting in NIR-IIc, we only performed modeling and calculated the PSF of confocal microscope in the NIR-IIc window as a function of the pinhole diameter. When a 25X objective (NA = 1.05) and a 1540-nm laser was used, and the fluorescence was collected in NIR-IIc window, the calculated PSF shows FWHMs of ~ 0.78 μm and ~ 1.72 μm in lateral and vertical direction, respectively (Supplementary Fig. 9).

### Data processing

The raw data of confocal microscopy was processed by a Gaussian filter in ImageJ. 3D rendering and multi-color fluorescence image merging were also performed in ImageJ or ImarisViewer 9.6.0 (Oxford Instruments). The FWHM was measured in Origin 9.0. The standard deviation and mean were calculated by Origin 9.0. The PSF was calculated in Matlab (R2019b).

### Mouse handling and tumor xenograft

Mouse handling was approved by Stanford University’s administrative panel on Laboratory Animal Care. All experiments were performed according to the National Institutes of Health Guide for the Care and Use of Laboratory Animals. BALB/c female mice were purchased from Charles River. Bedding, nesting material, food and water were provided. 3-week-old BALB/c mice (weight: ~ 7 g) were shaved using hair remover lotion (Nair, Softening Baby Oil). Mice were randomly selected from cages for all experiments. During *in vivo* imaging, all mice were anaesthetized by a rodent anesthesia machine with 2 l min^−1^ O_2_ gas mixed with 3 % isoflurane.

### In vivo wide-field NIR-II fluorescence imaging

The NIR-II wide-field fluorescence images were recorded by a 2D water cooled InGaAs camera (Ninox640, Raptor Photonics) working at −21 °C. For two-plex imaging, aMECA-79-QDc and p-FE were excited by an 808-nm continuous-wave diode laser. The fluorescence of aMECA-79-QDc and p-FE was collected in 1500-1700 nm and 1200-1400 nm, respectively. The actual excitation intensity was ~ 70 mW cm^−2^. The fluorescence signal was collected by two achromatic lenses to the camera after filtered by corresponding long-pass and short-pass filters.

### In vivo NIR-IIc confocal microscopy of mouse head

To image the blood vessels distribution in mouse head non-invasively (Fig. 3), a 3-week-old BALB/c mouse was intravenously injected with 200 μL NIR-IIb P^3^-QDb (O.D. = 5 at 808 nm) and 200 μL NIR-IIc P^3^-QDc (O.D. = 5 at 808 nm) sequentially through the tail vein. NIR-IIb and NIR-IIc confocal microscopy was performed 30 min after injection with a 25X objective (NA = 1.05). NIR-IIc confocal microscopy of intact mouse head was performed with a 5-μm scan increment along *z* direction. The integration time was linearly increased from 5 s to 30 s corresponding to the first frame at scalp surface and the last frame in the brain tissue. NIR-IIb and NIR-IIc fluorescence was collected in 1500-1700 nm and 1750-2000 nm windows, respectively. A 1319-nm and 1540-nm laser was used to excite NIR-IIb and NIR-IIc fluorescence, respectively.

### In vivo NIR-IIc confocal microscopy of inguinal lymph nodes

For two-color confocal microscopy of HEVs in iLN and blood vessels around the iLN, a 3-week-old BALB/c mouse was first injected with 200 μL aMECA-79-QDc (O.D. = 0.5 at 808 nm) intravenously. 24 h later, 200 μL p-FE (O.D. = 5 at 808 nm) was injected into the tail vein to label the blood vessels. 10 min post injection, two-color NIR-II confocal microscopy was performed. An 808-nm laser and a 1540-nm laser was used to excited p-FE and aMECA-79-QDc, and the fluorescence was collected in 1200-1400 nm and 1750-2000 nm, respectively. A 5X objective (NA = 0.12) or a 25X objective (NA = 1.05) was used. For 3D imaging, the scanning step was 5 μm and the integration time was linearly increased from 5 s to 20 s.

## Acknowledgments

This study was supported by the National Institutes of Health NIH DP1-NS-105737. We thank Kevin Taylor from JASCO who helped measuring the UV-Vis-NIR absorbance spectrum of water using their V-770 Spectrophotometer. F.R. thanks the Fonds de recherche du Québec – Nature et technologies (FRQNT) for funding. Single Quantum acknowledges support from the EIC SME Phase 2 project SQP, grant no. 848827.

## Author contributions

H.D. and F.W. conceived and designed the experiments. H.D. and F.W. designed the optical system. F.W. set up the optical system. F.W., F.R. and Z.M. performed the experiments. F.R. synthesize the PbS/CdS QD. R.G., I.E.Z., J.W.L., A.F. and J.Q.D. developed the SNSPD optimized in 1550-2000 nm window. F.W., F.R., Z.M., L.Q., C.X., A.B., J.L. and H.D. analyzed the data. F.W. and H.D. wrote the manuscript. All authors contributed to the general discussion and revision of the manuscript.

## Materials & Correspondence

Correspondence and requests for materials should be addressed to H.D.

## Supplementary Materials

### SUPPLEMENTARY VIDEO CAPTIONS

**Supplementary Video 1 |** Animated mouse brain video from non-invasive in vivo NIR-IIc confocal microscopy images of vasculatures in intact mouse head. The volumetric imaging was done using a 1540 nm-illumination for one-photon excitation and 1750-2000 nm detection for NIR-IIc P^3^-QDc circulating in brain vasculatures. The video here shows 3D images of intact mouse head with clearly resolved scalp, skull, meninges and brain cortex layers taken with 5-μm *z* increment in depth. The imaging time for each z-plane was linearly increased from 5 s to 30 s. A 25X objective (NA = 1.05) was used.

### SUPPLEMENTARY NOTE 1

#### Point spread function simulations of confocal microscope

The point spread function (PSF) of confocal microscope (PSF_confocal_) is given by

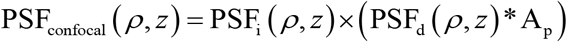

where PSF_i_ is the illumination PSF, PSF_d_ is the detection PSF, and A_p_ is the pinhole function. The * symbol represents convolution. *ρ* is the radius in (*x,y*) plane and *z* is the *z*-component of cartesian coordinates.

The illumination point spread function can be modeled by a Gaussian-Lorentzian profile^1^:

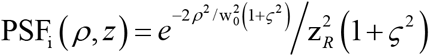

where 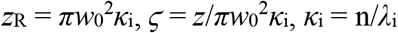, *w*_0_ is the waist of gaussian beam. *κ_i_* is the wavenumber. *λ_i_* is the excitation wavelength. n is the index of refraction of the surrounding medium.

PSF_d_ can be modeled by circular pupil as^1^:

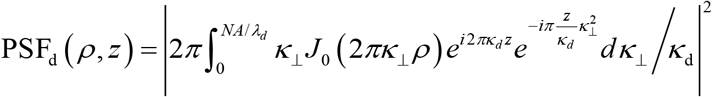

where NA is the numerical aperture of objective. *λ*_d_ is the emission wavelength. *κ*_d_ = n/*λ*_d_. *J*_0_ is the zeroth order cylindrical Bessel function.

The pinhole of confocal microscope can be defined by the unitless aperture function A_p_. For a circular aperture of radius *a_p_*, the Fourier transform of the pinhole function can be given as 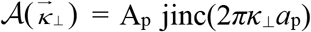, where 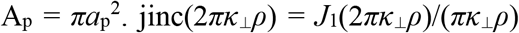, where *J*_1_ is the cylindrical Bessel function of order 1.

### SUPPLEMENTARY FIGURES

**Supplementary Figure 1.**
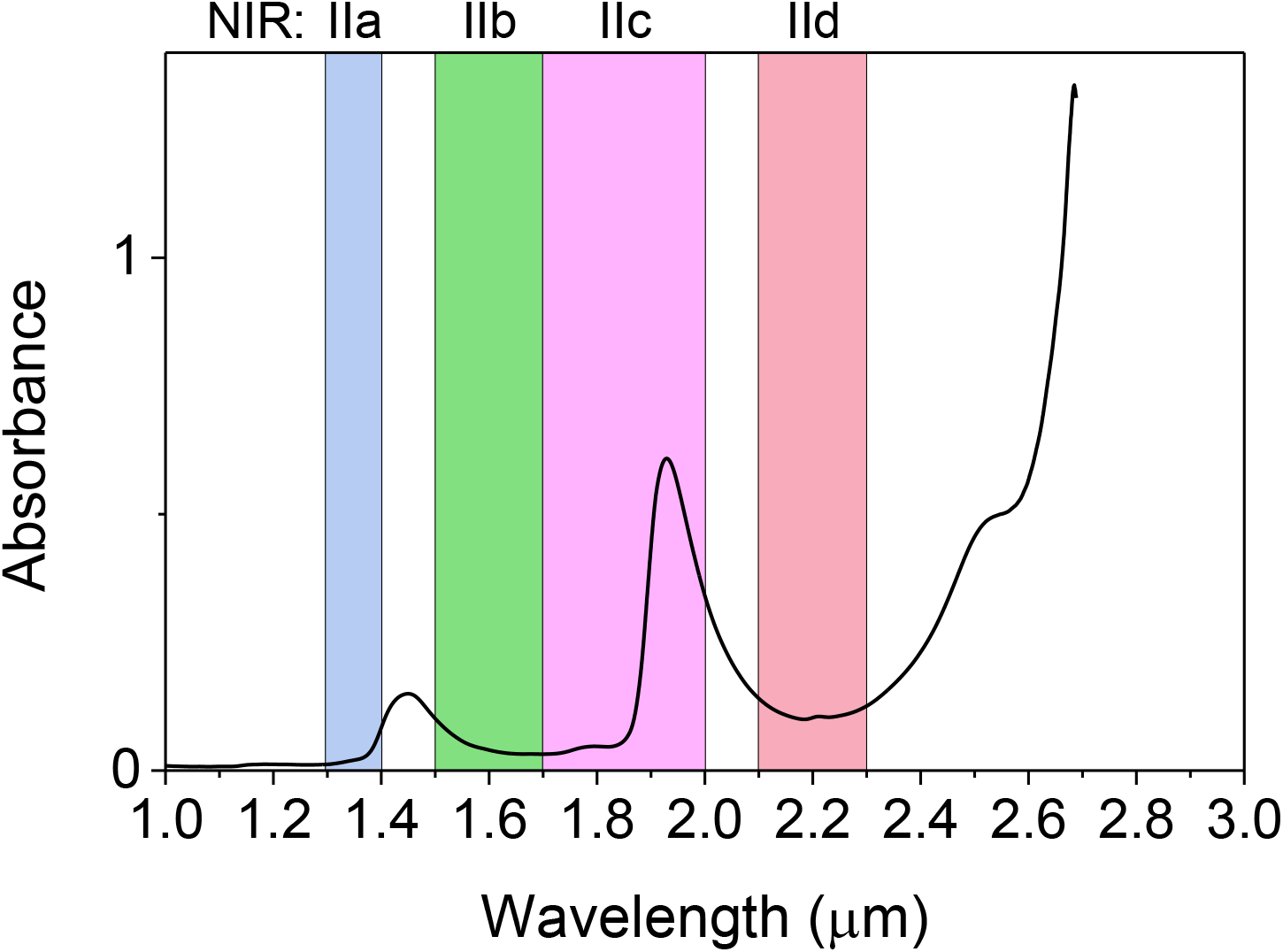
The absorbance of water measured in a cuvette with 0.1 mm light path. The UV-Vis-NIR absorbance spectrum of water was measured using V-770 Spectrophotometer (JASCO).

**Supplementary Figure 2.**
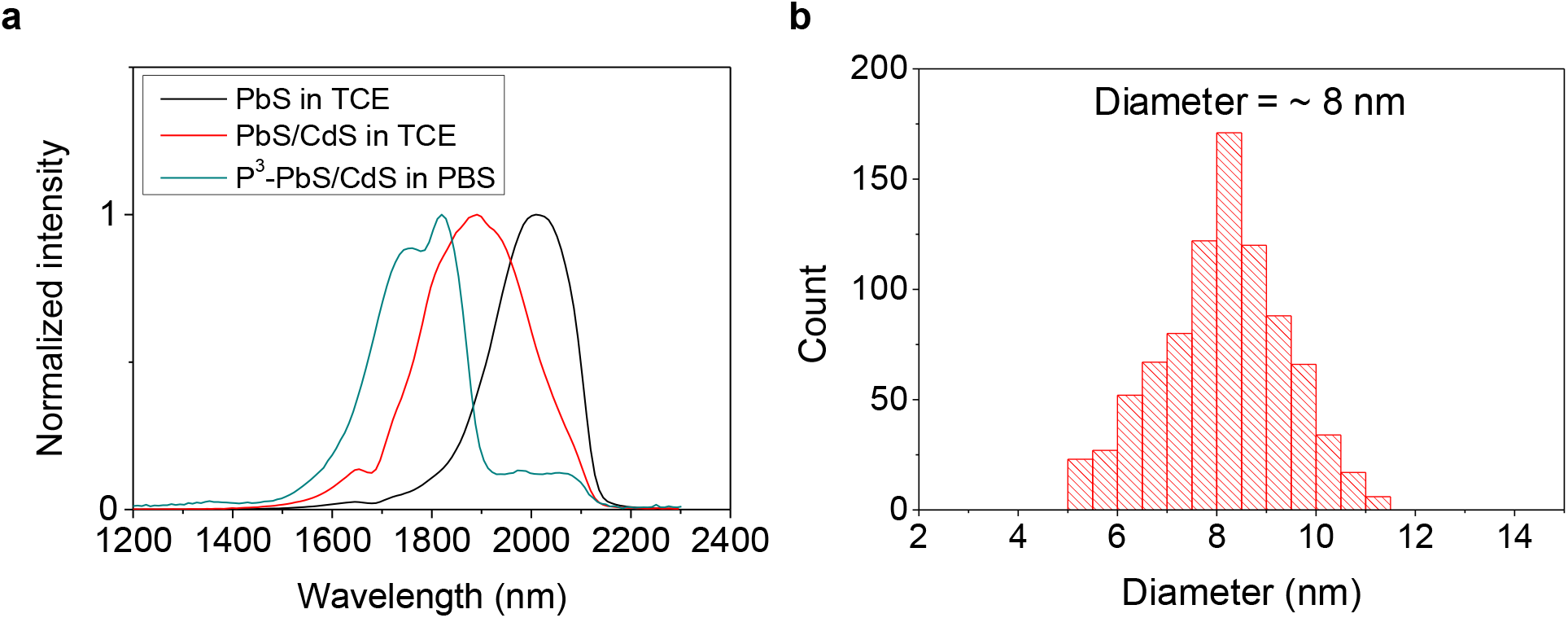
Characterization of NIR-IIc core/shell PbS/CdS quantum dots (named ‘QDc’ in this work). (**a**) Fluorescence emission spectrum of PbS core, core/shell PbS/CdS QD in tetrachloroethylene (TCE) and hydrophilic cross-linked P^3^ coated PbS/CdS QD in PBS after aqueous transfer. (**b**) Size-distribution histogram of QDc prior to P^3^-coating measured from TEM images.

**Supplementary Figure 3.**
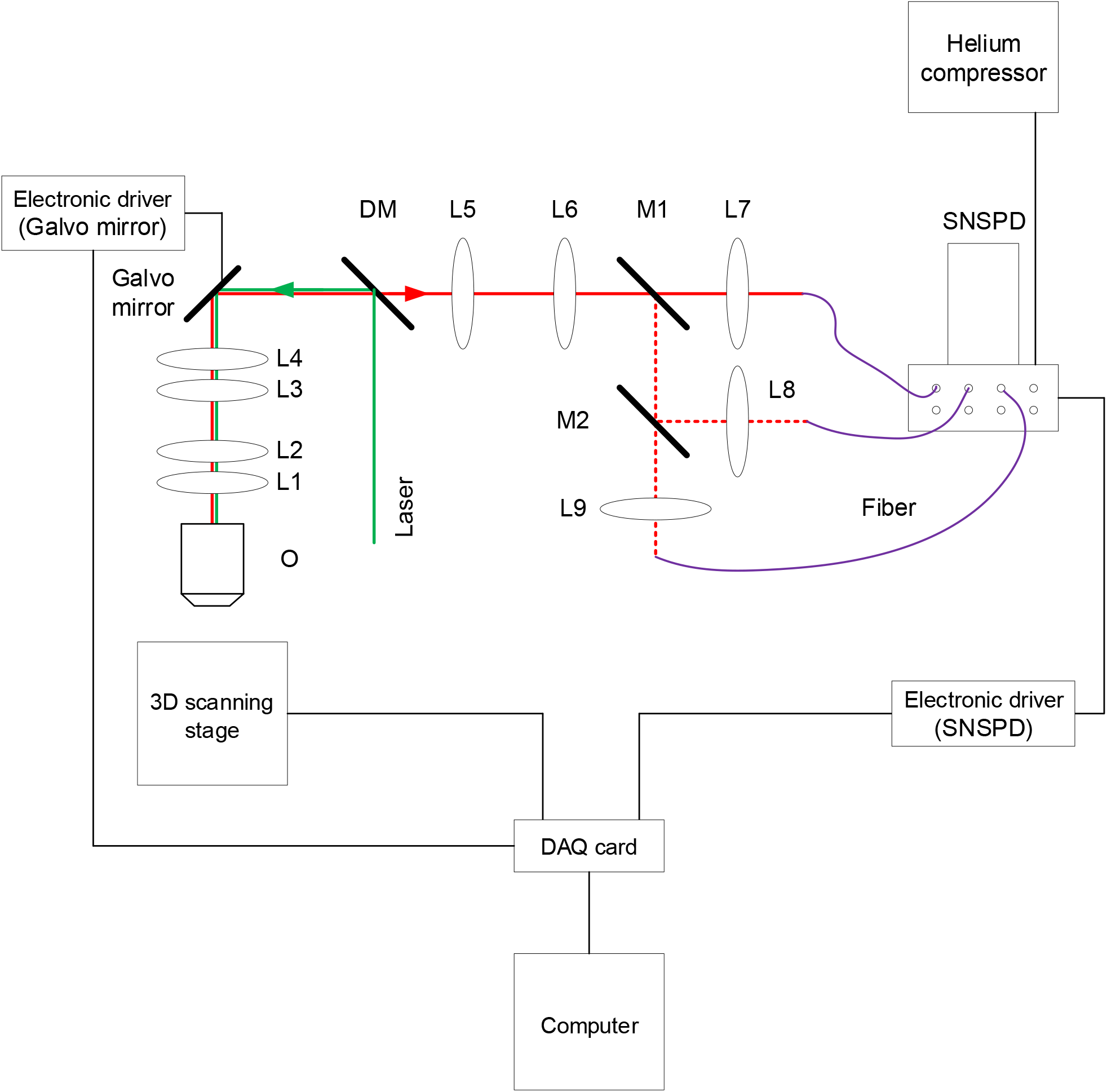
A schematic of the confocal microscopy in the second near-infrared window. The components are as follows: objective (O), achromatic lenses (L1 – L9), mirrors (M1, M2), dichroic mirror (DM). Three SNSPD detectors were selected by removable mirrors (M1, M2) A laser beam with wavelength of 808 nm, 1319 nm or 1540 nm was reflected by a galvo mirror (GVS002, Thorlabs) into a Plössl scan lens (constructed from two achromatic doublets, L3, L4, AC508-150-C, Thorlabs), a Plössl tube lens (constructed from two achromatic doublets, L1, L2, AC508-300-C, Thorlabs) and then focused by an objective into a sample (Supplementary Fig. 2). Fluorescence signal was collected by the same objective, tube lens and scan lens. After de-scanned by the galvo mirror and resized by two achromatic doublets (L5, AC254-100-C and L6, AC508-150-C), the fluorescence light was focused into three single mode (SM) fibers (P1-SMF28E-FC-5, Thorlabs) by achromatic doublets (L7-L9, AC254-030-C, Thorlabs) after filtered by selected emission filters. Then the SM fiber transmitted the fluorescence to NIR-IIb or NIR-IIc SNSPD.

**Supplementary Figure 4.**
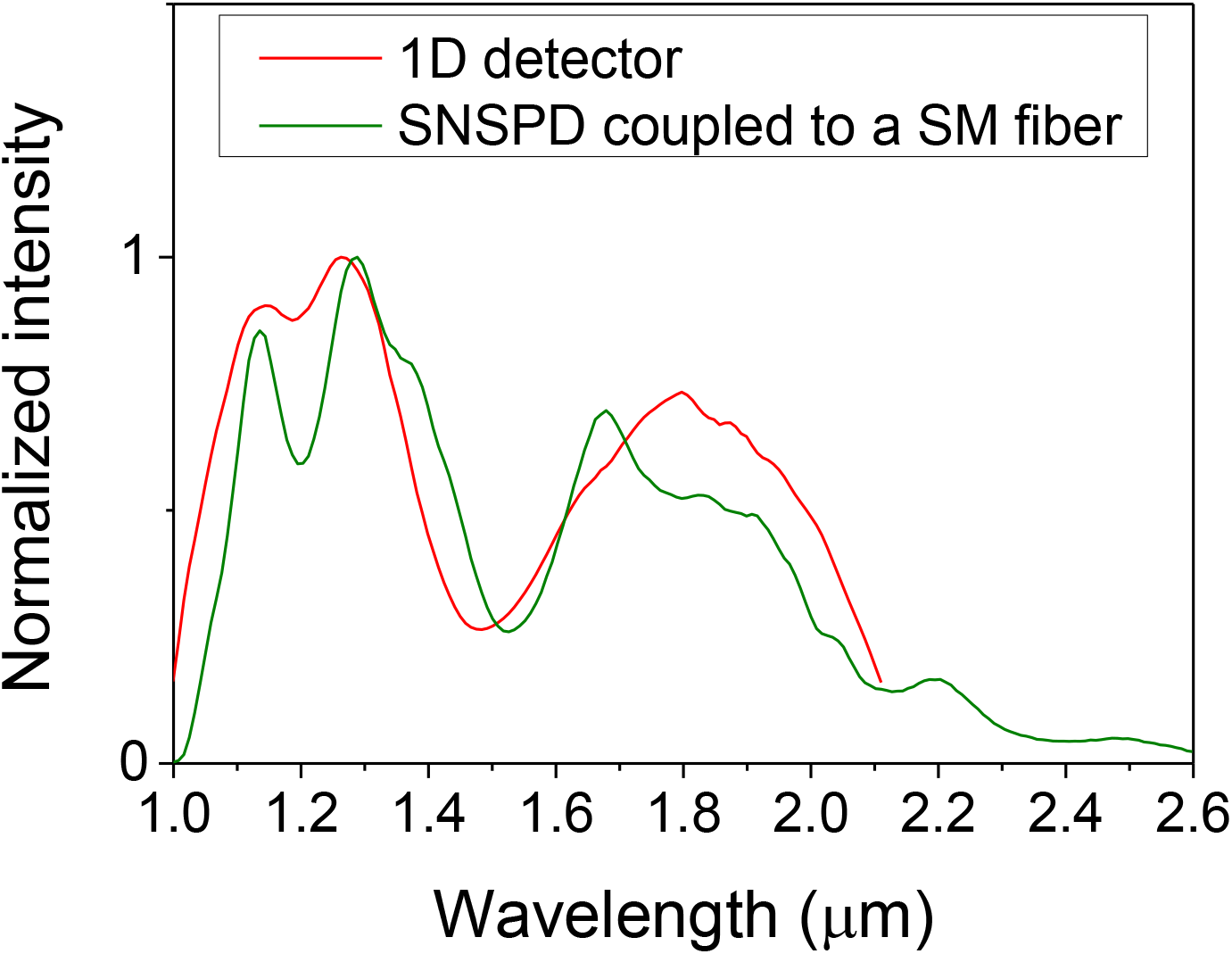
Single-mode (SM) fiber used for NIR-IIc confocal microscopy using the SNSPD. We mixed three PbS/CdS QDs of different average sizes to a suspension with emission covering the ~ 1000-2100 nm range. The overall emission spectrum was first measured by a commercial spectrometer with a 1D extended InGaAs camera (Princeton Instruments) without SM fiber coupling as shown by the red curve. The fluorescence was collected by a couple of lenses into the spectrometer. Then the spectrum of the mixed PbS/CdS QDs was measured again using the same spectrometer but connected to SNSPD coupled to a SM fiber (green curve). An 808 nm laser was used to excite the mixed QDs. The fiber used here was P1-SMF28E-FC-5 from Thorlabs with a specification of transmission in the 1260-1625 nm range. Our data here showed that the fiber also allowed transmission in the 1700-2000 nm (NIR-IIc), which was not described in the product datasheet. We tried various fibers and this fiber turned out to be optimal for NIR-IIc confocal imaging using SNSPD.

**Supplementary Figure 5.**
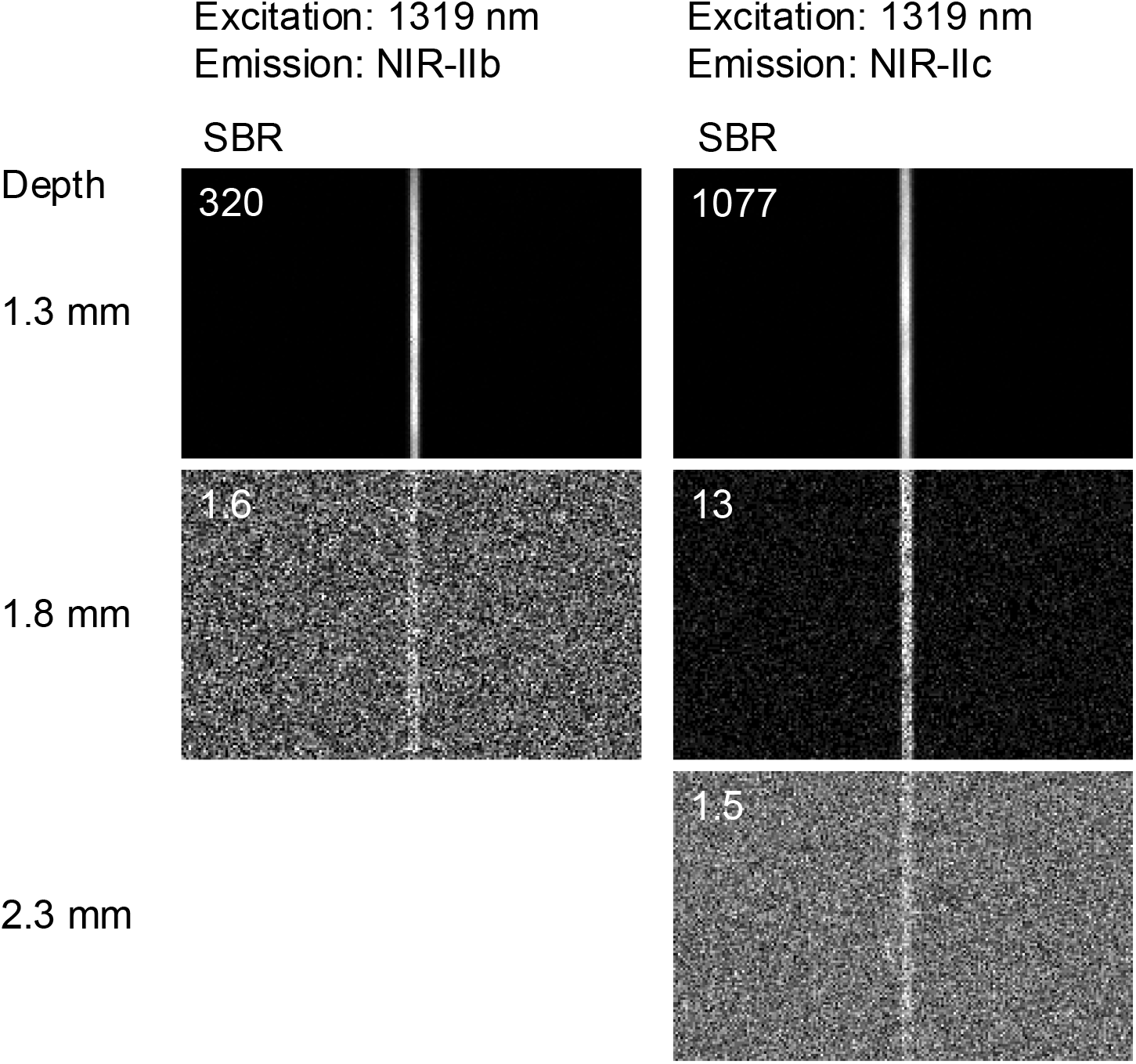
Imaging a capillary tube containing QDb or QDc immersed in 5% intralipid water solution. Fluorescence confocal images of a 50-μm diameter capillary tube filled with NIR-IIb QDb (left column) or NIR-IIc QDc (right column) immersed at different depths in 5% intralipid acquired with a confocal microscope equipped with SNSPD. A 1319-nm laser and a 5X objective (NA = 0.12) were used for NIR-IIb and NIR-IIc confocal microscopy. NIR-IIb and NIR-IIc fluorescence was collected in 1580-1700 nm or 1750-2000 nm windows, respectively. At 1.3 mm depth, imaging under 1319 nm laser excitation afforded higher SBR than NIR-IIc imaging under 1540 nm laser excitation, but with a faster SBR decrease at larger depth. The maximum imaging depth decreased from ~ 2.3 mm and ~ 2.6 mm to ~ 1.8 mm and ~ 2.3 mm for NIR-IIb and NIR-IIc confocal microscopy using SNSPD respectively. These results confirmed that longer emission wavelength in NIR-IIc allowed for deeper imaging penetration depth in scattering media than in NIR-IIb.

**Supplementary Figure 6.**
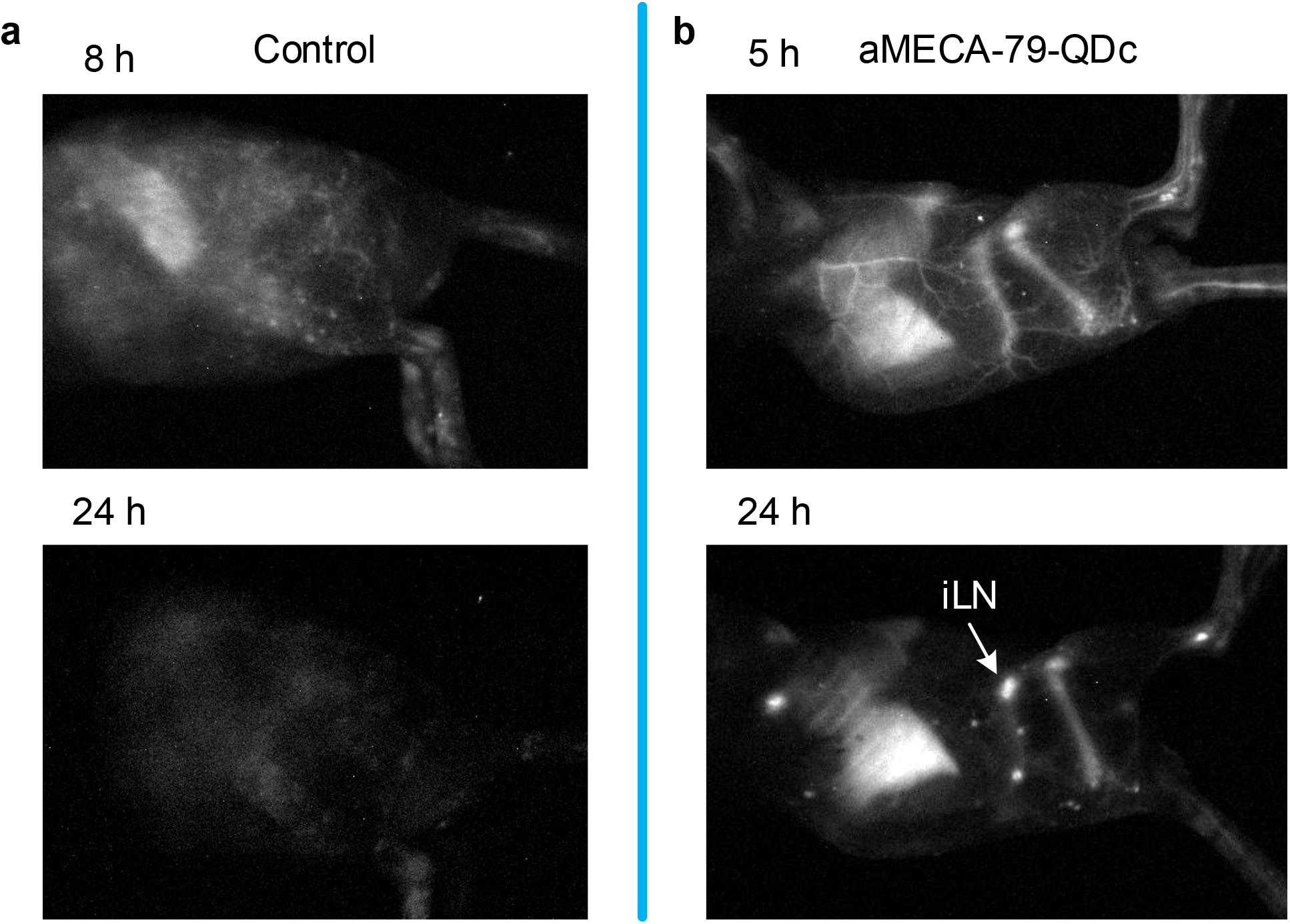
Wide-field images of a Balb/c mouse recorded after intravenous injection of (**a**) free P^3^-QDc and (**b**) aMECA-79-QDc at two different timepoints for each case. The fluorescence was collected in 1500-1700 nm under an 808 nm laser excitation. In (**a**) we did not observe apparent fluorescence signal in the inguinal lymph node when free QDc were injected. In contrast, in (**b**) the inguinal lymph node showed strong signal 24 h post injection of aMECA-79-QDc intravenously, suggesting targeted iLN HEVs labeling by QDc conjugated with anti-MECA-79.

**Supplementary Figure 7.**
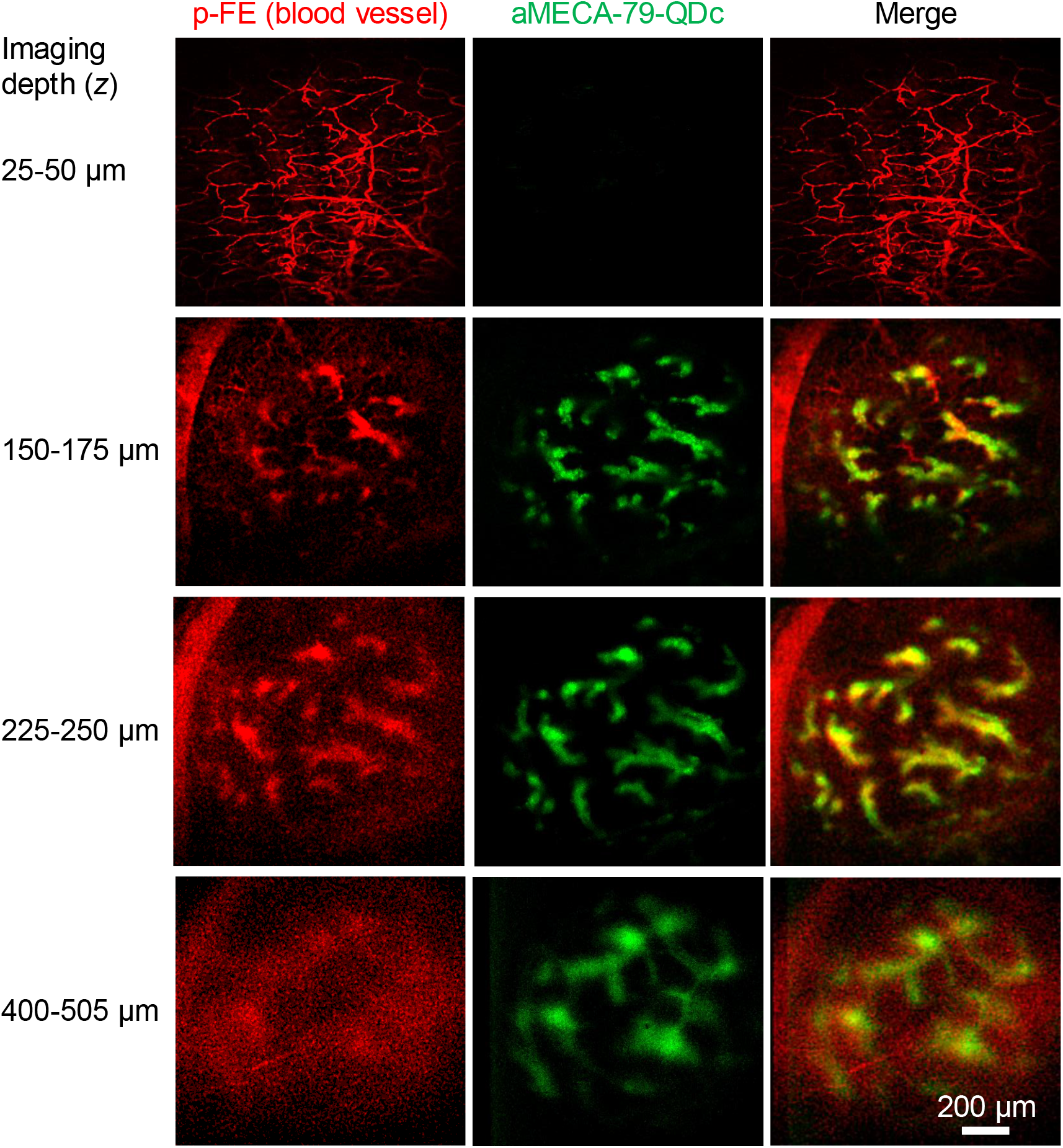
Non-invasive two-color *in vivo* NIR-II confocal microscopy imaging of inguinal lymph nodes in mice at various depths under the skin. High-resolution confocal microscopy of blood vessels (red color, vessels contained circulating p-FE dye; excitation: 808 nm, emission: 1200-1400 nm) and HEVs in iLN were targeted and labelled specifically by aMECA-79-QDc (green, excitation: 1540 nm, emission: 1750-2000 nm). Imaging was done with a 5 μm scan increment along *z* direction. The imaging time per plane was linearly increased from 5 s to 15 s from the first frame at the skin surface to the last plane inside the iLN. Imaging depths are labeled at the left of the figure. Notice that all vessels contained the circulating p-FE dye, and only HEVs are specifically targeted and labeled by the antibody-QDc conjugate.

**Supplementary Figure 8.**
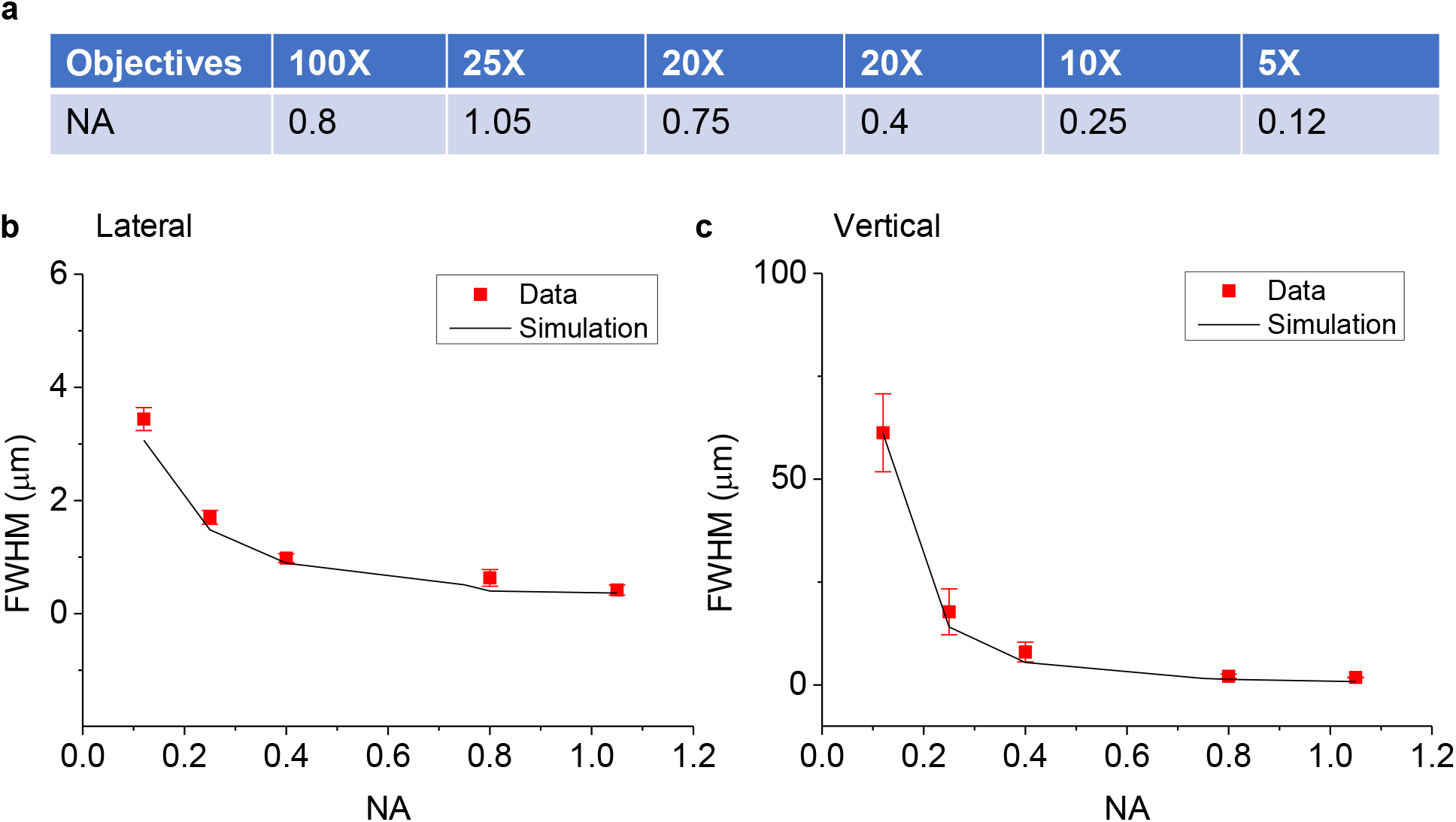
Resolution of confocal microscopy with SNSPD analyzed by experimental measurements and theoretical calculations. (**a**) Objectives with different magnifications and numerical apertures were used in this analysis. Comparison of experimentally measured FWHMs and FWHMs of theoretically calculated PSFs in (**b**) lateral or (**c**) vertical directions. For experimental measurements, we imaged 300 nm polystyrene beads containing a NIR-II organic dye (p-FE)^2^ (emission mostly in 1000-1300 nm range) with different objectives listed in (**a**) by the confocal microscope with SNSPD coupled to a single mode fiber with ~ 10-μm mode field diameter. Fluorescence emission > 1100 nm was collected and 658-nm laser was used. The FWHMs of nanoparticles in the lateral and vertical directions were measured from the confocal microscopy images. For calculation, the model in Supplementary Note 1 was used to calculate the PSF based on parameters the same to the experimental conditions. The FWHMs of PSFs were measured to compare with experimental measured FWHMs of beads with excellent agreement.

**Supplementary Figure 9.**
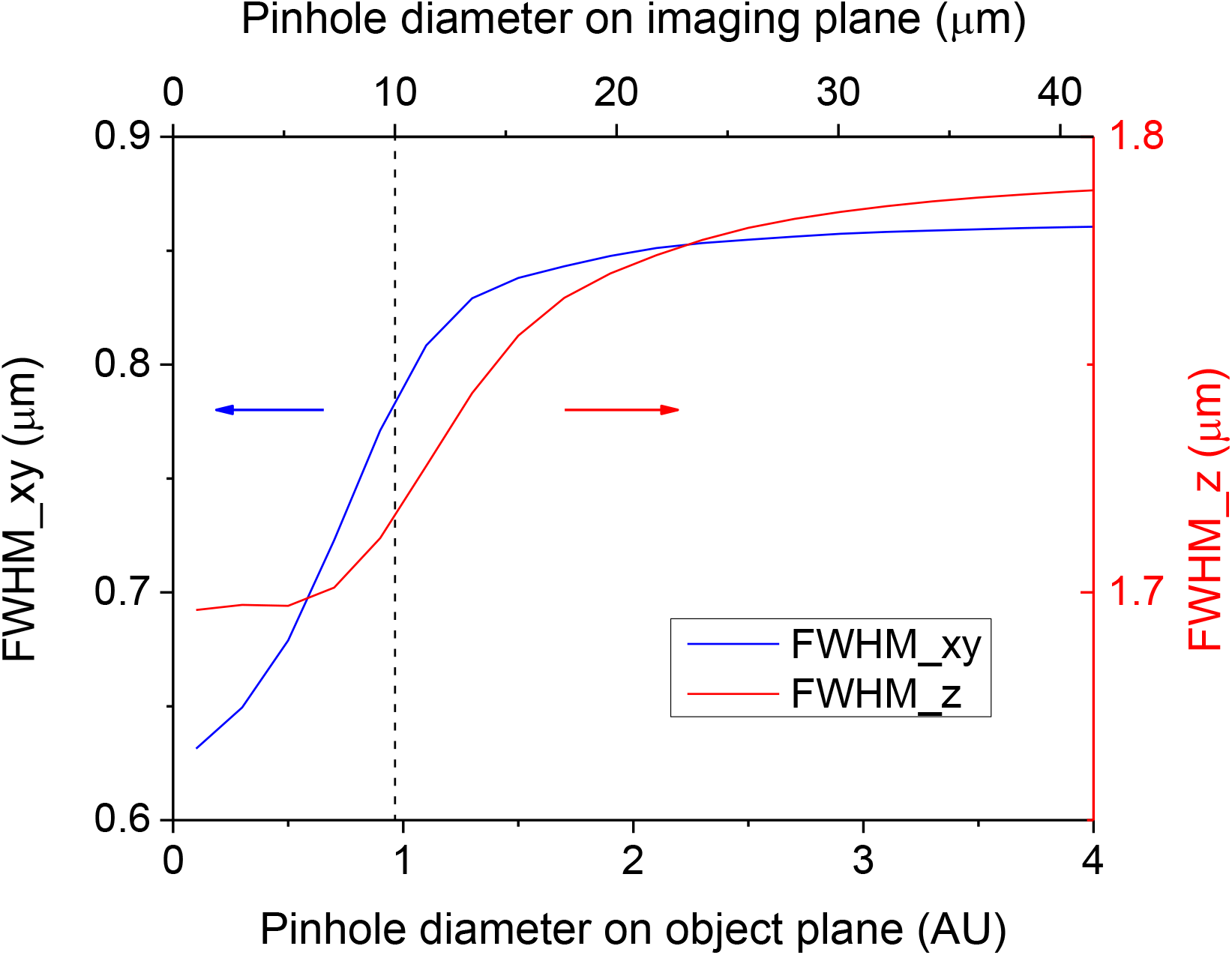
The influence of the pinhole diameter on the resolution of NIR-IIc confocal microscopy. In this simulation, the excitation was 1540 nm and fluorescence emission was ~ 1800 nm. A 25X objective (NA = 1.05) was used. The pinhole diameter was normalized as Airy units, 1 AU = 1.21*λ*_d_/NA_d_, where NA_d_ is the numerical aperture of objective. *λ*_d_ is the emission wavelength. A single mode fiber with ~ 10-μm mode field diameter (corresponding to ~ 0.96 AU, dash line) was used in our experiments and worked as a pinhole. This combination enables PSF with FWHMs of ~ 0.78 μm and ~ 1.72 μm in lateral and vertical direction, respectively

